# Effect of stimulation parameters for transcranial temporal interference stimulation to enhance occipital alpha-oscillations in healthy participants

**DOI:** 10.64898/2026.09.14.751477

**Authors:** Johannes Stalter, Heiko Stecher, André Aleman, Christoph S. Herrmann, Karsten Witt

**Affiliations:** Department of Neurology, Carl von Ossietzky Universität Oldenburg, 26122 Oldenburg, Germany; University Hospital of Neurology, Evangelical Hospital Oldenburg, 26122 Oldenburg, Germany; University of Groningen, Cognitive Neuroscience Center, University Medical Center Groningen, 9713 GZ Groningen, The Netherlands; Experimental Psychology Lab, Carl von Ossietzky Universität Oldenburg, 26129 Oldenburg, Germany; Faculty of Psychology and Neuroscience, Maastricht University, Maastricht, The Netherlands; Center of Neurosensory Sciences, Carl von Ossietzky Universität Oldenburg, 26129 Oldenburg, Germany; Cluster of Excellence “Hearing4all”, Carl von Ossietzky Universität Oldenburg, 26129 Oldenburg, Germany

## Abstract

Transcranial temporal interference stimulation (tTIS) has recently emerged as a non-invasive method for modulating neural activity using interfering high-frequency electric fields. Although behavioral and neuroimaging effects of tTIS have been reported, evidence for electrophysiological modulation remains inconsistent, and it is unclear whether observed effects are attributable to temporal interference itself or to other components of the signal, e.g. high-frequency carrier signals. We therefore investigated whether tTIS at to the individual alpha frequency (IAF) modulates occipital alpha power compared with both sham stimulation and an active high-frequency carrier control.

Eighteen healthy participants were enrolled in a randomized, double-blind, within-subject crossover study; 17 participants were included in the final analysis. Each participant completed three stimulation sessions consisting of three conditions (tTIS, carrier control, sham stimulation). tTIS was delivered for 20 minutes using two high-frequency signals at 1000 Hz and 1000 Hz + IAF, whereas the carrier control used two 1000-Hz signals without an envelope modulation. EEG was recorded before and after stimulation during a visual vigilance task. Individual electric-field distributions in the occipital cortex were estimated using structural MRI-based finite-element simulations.

A significant main effect of stimulation condition on the change in occipital alpha power was observed (F(2,32)=5.69, p=.008, η^2^p=.26). Post-hoc comparisons showed a significantly greater increase in alpha power following tTIS compared with both the carrier control (mean difference=0.30, adjusted p=.036, Cohen’s d=0.686) and sham stimulation (mean difference=0.31, adjusted p=.037, Cohen’s d=0.683). Carrier and sham conditions did not differ (adjusted p=1.000). No behavioral effects were observed. The magnitude of the alpha-power change was not significantly associated with simulated electric-field strength in the occipital target region. tTIS was well tolerated, with no stimulation-related study discontinuations.

tTIS targeting the occipital cortex therefore produced significant post-stimulation enhancement of alpha power beyond both sham and high-frequency carrier stimulation. The absence of an effect in the carrier condition supports the interpretation that the electrophysiological aftereffect depends on temporal interference rather than high-frequency stimulation components alone. These findings provide evidence for frequency-specific modulation of cortical oscillations by tTIS and support further investigation of its underlying mechanisms and stimulation parameters.

## Introduction

Over the last decades, deep brain stimulation has emerged to a crucial technique to modulate brain oscillations and functions of targets in deep brain regions. It is nowadays used as a treatment option for several neurological and psychiatric diseases.^1–5^ However, it is still an invasive procedure with several risks side effects, even after completed operation.^6^ With its first mention as a non-invasive deep brain stimulation method, transcranial temporal interference stimulation (tTIS) has been recently pushed as a method to reach deep brain targets with only little off-target stimulation and only few side effects.^7,8^ For this technique, at least two independent high-frequency electrical fields with a small frequency difference (e.g., 2 and 2.1 kHz; carrier frequencies) are applied transcranially. The superposition of these fields results in an envelope modulation at the difference frequency (in this example, 0.1 kHz), in deeper brain regions (region of interest; ROI) which can be targeted by optimization of the stimulation parameters. This reduces simultaneous stimulation of superficial cortical areas and thus overcomes the limited depth focality of conventional electro-(magnetic) stimulation methods. This envelope modulation can then influence neuronal activity in the target region. For an overview of this technique and its possible application, see Stalter et al. 2025.^9^ Since its introduction, multiple studies have proven its efficiency in modulating behavior and (electro)-physiological outcome measures like local field potential (LFP) and functional magnetic resonance imaging (fMRI) responses in healthy humans and patients.^10–14^ Hippocampal targeting enhanced performance in a face–name association task, whereas tTIS targeting the putamen improved motor-sequence performance and altered the BOLD response in the targeted region. In disease models like Parkinson’s disease, tTIS was able to improve clinical motor symptoms and to change beta activity in the basal ganglia in multiple studies.^10,15–17^ While these findings provide a solid foundation to investigate the behavioral outcomes and the changes in fMRI, studies investigating its effects on electroencephalography (EEG) are more heterogenous and show no clear trend. Von Conta et al. compared tTIS to transcranial alternating current stimulation (tACS) and a high frequency control condition and found no evidence for a significant difference between these three conditions for a change in occipital alpha power. This could be either interpreted as no effect of tTIS or as an effect of both, tTIS and control stimulation.^18^ Another study examined the effects of event related desynchronization (ERD), resting state alpha power and the behavioral outcome in a young and healthy group in comparison to tACS. While ERD increased, no change in alpha power was detected between the pre- and poststimulation data. Therefore, there are still open questions about the EEG effects of tTIS which could answer questions about the underlying mechanisms of this method. In the present study, we investigated the effects of tTIS on occipital alpha power in young and healthy participants to provide further evidence of the mechanism of tTIS. The target region of the occipital cortex was chosen to include as little as possible confounding factors which would have not been possible with deep brain regions, as for these technical key components like the actual electrical field strength are not exactly measurable.

## Methods

### Participants

Due to the uncertainties of the effect sizes with regard to the target site and the outcome, the estimation of the sample size is taken from a pre-study, involving transcranial temporal interference stimulation. 18 young, healthy participants with no known history of neurological or psychiatric disease, no reported intake of central-acting drugs or contraindications for MRI or transcranial electrical stimulation were recruited. The study was approved by the local ethical committee (2024-156) and was conducted in accordance with the Declaration of Helsinki.^19^ A pre-registration on OSF was created prior to the beginning of the data acquisition (10.17605/OSF.IO/WM9TG).

### Anatomical MRI

To simulate the electrical stimulation, structural MRI images were obtained from all participants. The images were acquired at the Neuroimaging Unit at the Carl von Ossietzky University Oldenburg using a Siemens Magnetom Prisma 3T MRI scanner (Siemens, Erlangen, Germany). A T1-weighted sequence (TR = 2300 ms; TE = 4.16 ms; voxel size 1x1x1 mm) and a T2-weighted sequence (TR = 5000 ms; TE = 390 ms; voxel size 1x1x1 mm) were measured.

### EEG

The EEG-setup consisted of 28 active electrodes placed according to the international 10-10 system in an actiCAP (Brainproducts, Gilching, Germany). The EEG-signal was recorded using an actiCHamp amplifier (BrainProducts, Gilching, Germany). The reference electrode was placed on the tip of the nose while impedances were kept below 10 kW. A 3 minute pre-recording was used to define the individual alpha frequency (IAF) which was then used for the stimulation block.

### Simulation

To simulate stimulation-effects, SimNIBS 4.6 was used.^20^ Individual scans (T1 and T2) were segmented, and conductivity of tissue type was assigned to create individual meshes. Using these meshes, simulation on a single-subject level was computed. Simulations were run with an injected current of 2 mA peak-to-peak for electrode pairs positioned at C3/O1 and C4/O2. Using the melb-atlas, we simulated the electrical field in the occipital lobe adapted to the individual neuroanatomy in the subject space of each participant (Figure 4B).^21^ Simulation was performed post-hoc only and therefore had no influence on individual stimulation parameters. To test the correlation between the e-field and the EEG data, we calculated the difference between the change during sham and actual stimulation. This difference was then correlated with the simulation. We chose this approach to take the change due to the task or structural influences into account.

### Electrical Stimulation

The stimulation was applied by two pairs of electrodes placed at C3/O1 and C4/O2 (according to the international 10-10 system) attached to the scalp using ten20-paste (Weaver & Co, Aurora, CO, USA) with impedance below 10 kΩ. For stimulation, 5x5cm rubber electrodes were used with a current-intensity of 2 mA (peak-to-peak) since this is below the recommended maximum current density in skin of 0.1 mA/cm^2^ (= 2.5 mA zero-to-peak for 25cm^2^ electrodes). Two minimize potential side-effects, a ramp up and ramp down of 30 seconds was applied for all stimulation conditions. Three different conditions were chosen for this study. The verum stimulation consisted of two high-frequency signals both oscillating continuously at 1000 Hz and 1000 Hz + IAF. The sham condition was defined as two signals with a ramp up/down with no active signal in between. Lastly, the carrier condition was applied as two signals both at 1000 Hz, therefore no active envelope frequency was applied during this condition. Each condition lasted for 20 minutes, and the conditions were randomized for each participant. The order of the conditions was blinded to the participant and the examiner.

All signals were computed with Matlab (MATLAB, Natick, Massachusetts: The MathWorks Inc.) and sent to a digital-to-analog converter (Ni-USB 6251, National Instruments, Austin, TX, USA). The currents were delivered by two battery-driven, galvanically isolated constant current stimulators (Advanced DC Stimulator Plus, Neuroconn, Ilmenau, Germany).

### Task Design

A white fixation cross was presented on a grey screen during the task. The cross rotated at random intervals by 45° for 500 ms and participants were asked to press a button every time a rotation took place. The task was presented using Matlab (MATLAB, Natick, Massachusetts: The MathWorks Inc.) and PsychToolbox 3.^22,23^

### Experimental Design

After checking the in- and exclusion criteria, for each participant an MRI scan was obtained. The following three sessions all followed the same procedure with only the condition itself being different. After setting up the EEG and the stimulation equipment, each session started with a three minutes resting state EEG measurement to define the IAF. For this the EEG signal underwent individual component analysis (ICA) and the IAF was searched for by a peak detection in the alpha band (8 – 12 Hz). After this, the pre-measurement took place. No stimulation was applied during that time, and participants underwent a vigilance task. After this block, the intervention started and the vigilance task was performed again. This was followed by the post block where again only EEG was measured during the vigilance task. After the last block, the participants stayed in the lab for approximately 30 minutes to observe if any adverse side-effects occur.

### Data Analysis

The data was analyzed using Matlab (R2025a, Natick, Massachusetts: The MathWorks Inc.) and the Fieldtrip toolbox.^24^

#### IAF

The resting state EEG data were down sampled to 250 Hz to achieve faster processing. After demeaning and detrending, bad channels were rejected by visual approach or if noted down as bad during the recoding. An ICA was performed to remove eye-movement-artifacts manually based on a structured approach. Following this, a Fast-Fourier-Transformation was performed to examine frequency power between 1 and 20 Hz. IAF was defined as the highest peak between 7 and 13 Hz using MATLAB’s findpeaks()-function (with ‘NPeaks’=2, Sort=‘desc’). This frequency was then used as the IAF for stimulation.

#### EEG

The EEG data first was demeaned, detrended and filtered between 1 and 48Hz. After this, the data was cut into three blocks (pre/during/post). The continuous data for the pre and the post-block were segmented into non-overlapping 2-s epochs. Epochs containing gross artifacts were rejected using FieldTrip’s threshold-based artifact detection with a peak-to-peak range threshold of 500 [µV]. Power spectra were estimated for artifact-free epochs using a fast Fourier transform (mtmfft) with a Hanning taper. Data were zero-padded to 10 s, and spectral estimates were obtained from 1 to 30 Hz in 0.1-Hz steps. Power was subsequently averaged across epochs and then over all parietal channels. The averaged parietal power at stimulation frequency was then used for the further analysis.

### Statistics

Statistical analysis was calculated with Matlab (R2025a, Natick, Massachusetts: The MathWorks Inc.). The pre-post change in relative alpha-power was evaluated in a 2x3 repeated measurement ANOVA with a 2-level factor block (pre/post) and a 3-level factor condition (tTIS/Carrier control/sham) with the relative change in alpha power as the depended variable. A repeated measurement ANOVA was also used to analyze the behavioral outcome (reaction time and correctness). A significance level of 0.05 was used for all analyses; post-hoc tests were correct using Bonferroni correction.

## Results

### Demographical Characteristics and side effects

Based on a previous study with a similar design, we estimated a sample size of 16 participants.^25^ To account for potential dropouts, we enrolled 18 participants of which only one had to be excluded for technical reasons, i.e. missing EEG recording in one block. The analyzed group consisted of 17 participants with a mean age of 27±3.2 with a gender distribution of ten females and 7 males. For this study, we completed a total of 51 stimulation sessions. The most common side-effects were discomfort under the electrodes, tiredness, concentration difficulties and itching at the side of the electrodes. However, it is noteworthy that most participants wrote the length of the experiment as the probable cause of the tiredness. Figure 1 shows the frequency of side-effects and the reported severeness on a scale 0-3 (0 = no reported side-effect, 3 strong sensations of the side-effect). No stimulation session was discontinued due to unpleasant sensations, and no study drop out was reported due to the stimulation.

**Figure 1.**
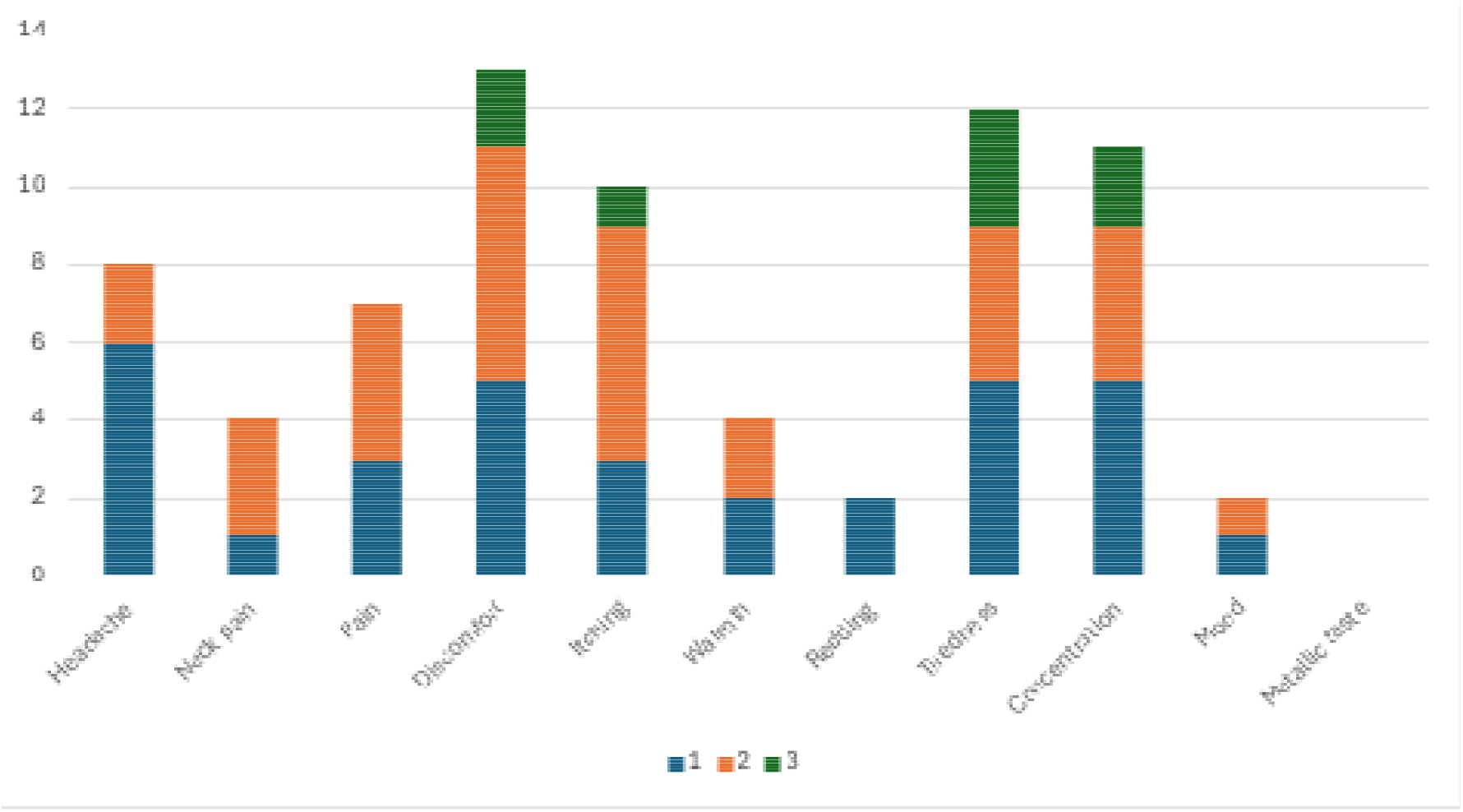
Frequency of reported side-effects and how severe they were experienced. For this study a total of 51 sessions were completed.

### Task results

The mean reaction time for tTIS was 0.563 seconds (SD=0.012), for the carrier control 0.652 seconds (SD=0.288) and for the sham condition 0.591 seconds (SD=0.177). Comparing the reaction times during the task between different conditions, a repeated measurement ANOVA revealed no significant effect of stimulation (F(2,32) = 2.51, p = .097, ph^2^ = .138) (figure 2).

**Figure 2.**
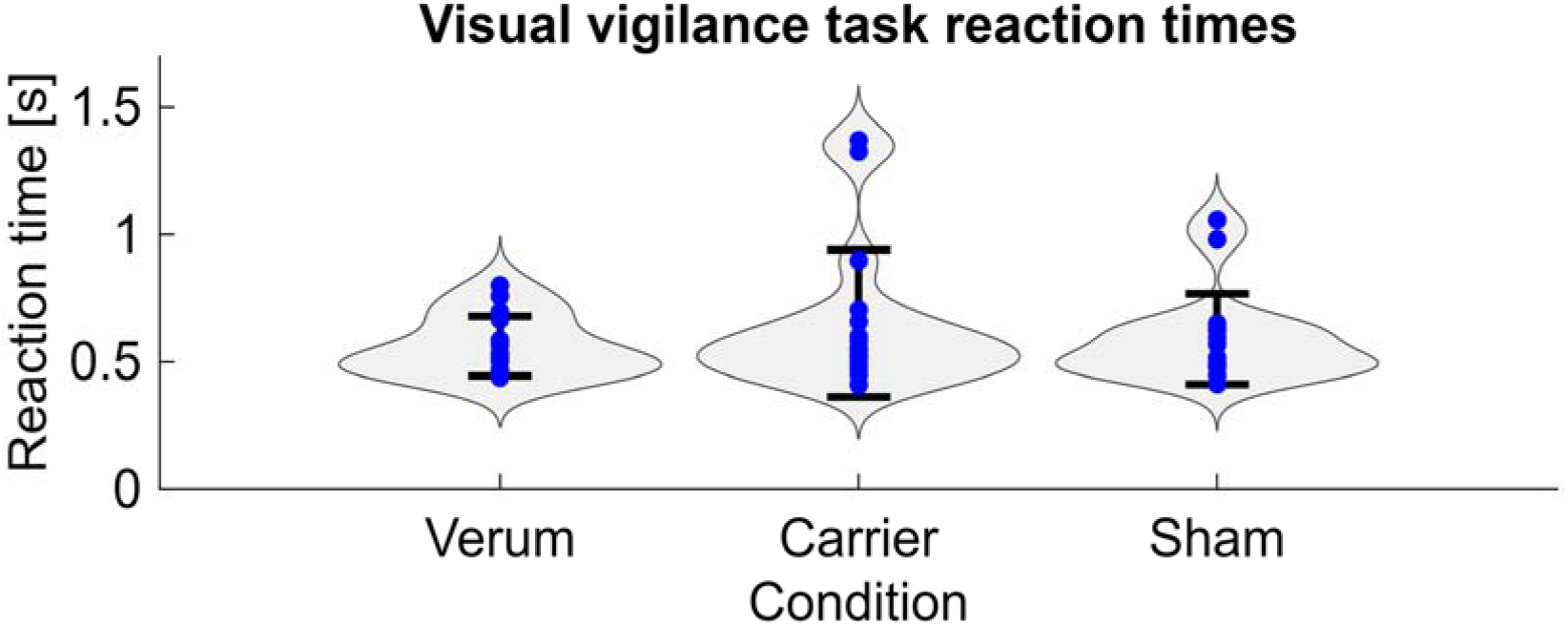
Reaction times measured during the visual vigilance task for all three conditions. No significant effect on the reaction was found for the different conditions. Blue dots represent individual data points.

### Parietal alpha-power

The repeated-measures ANOVA revealed a significant main effect of Stimulation (*F*(2, 32) = 5.69, *p* = .008, ph^2^ = .26). Bonferroni-adjusted pairwise post-hoc comparisons showed that the alpha power increase was significantly higher in the tTIS condition than in both the high frequency carrier condition (mean difference = 0.30, SE = 0.10, adjusted *p* = .036, 95% CI 0.02 - 0.58, Cohen’s d = .686) and the sham condition (mean difference = 0.31, SE = 0.11, adjusted *p* = .037, 95% CI 0.02 - 0.60 Cohen’s d = .683). There was no significant difference between the Carrier and Sham conditions (mean difference = 0.01, SE = 0.10, adjusted *p* = 1.000, 95% CI −0.25 - 0.26, Cohen’s d = -.023). Figure 3 depicts the relative change in alpha band power in the three conditions.

**Figure 3.**
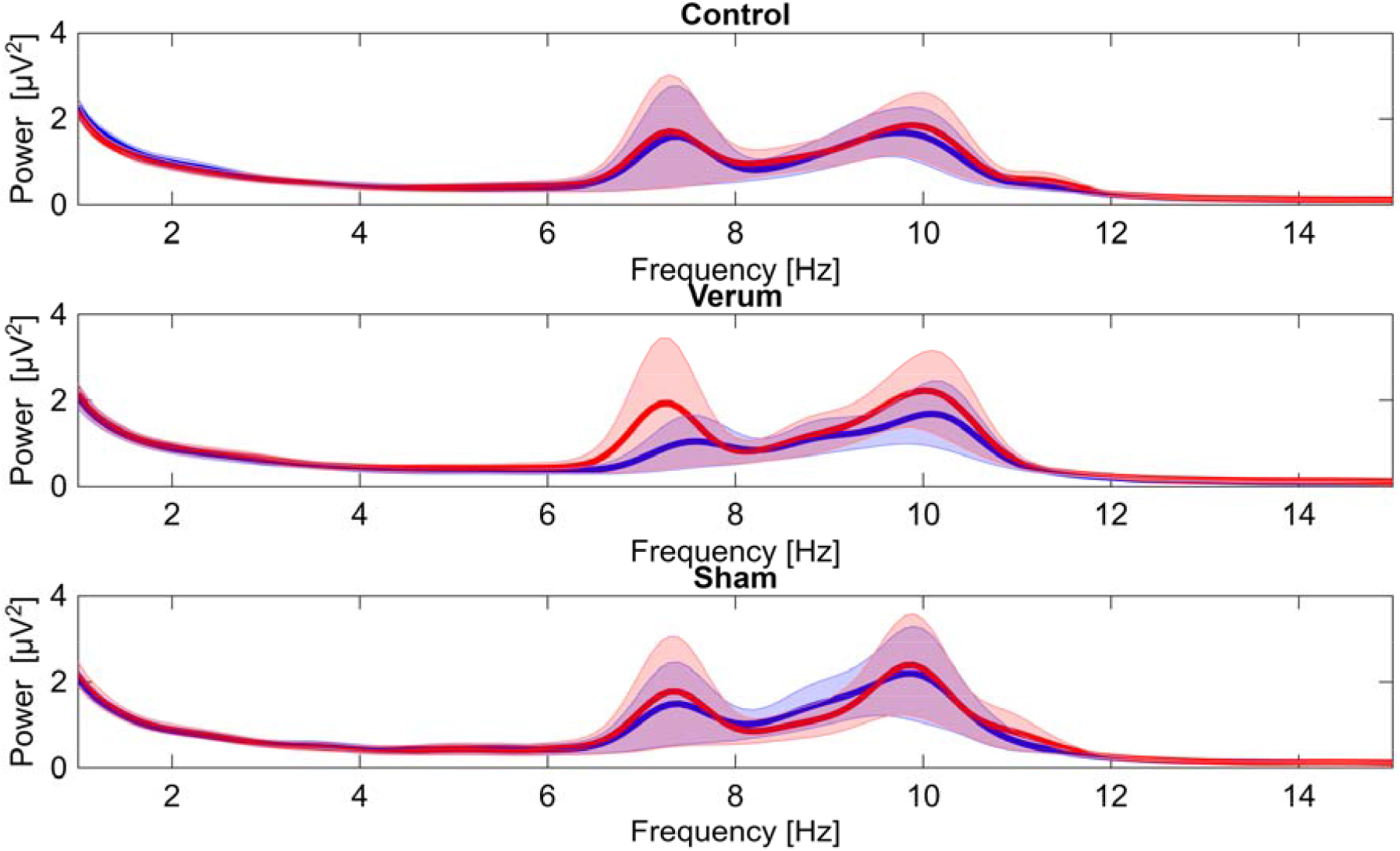
Upper Panel) Change in alpha power in the control condition for the pre and the post measurement with no significant difference between pre/post. Middle panel) Same presentation for the tTIS (verum) condition showing the significant difference in the alpha band. Lower panel) Presentation of the change during the sham condition again with no significant difference. Note that the peak at roughly 7.5 Hz was due to a single subject with high amplitudes.

### E-field correlation

The calculated difference between the relative alpha band power between pre and post measurement was correlated with the simulated e-field in the occipital lobe. There was no significant correlation between both variables evident (rho = .268, p = .298, Pearson correlation) (figure 4).

**Figure 4.**
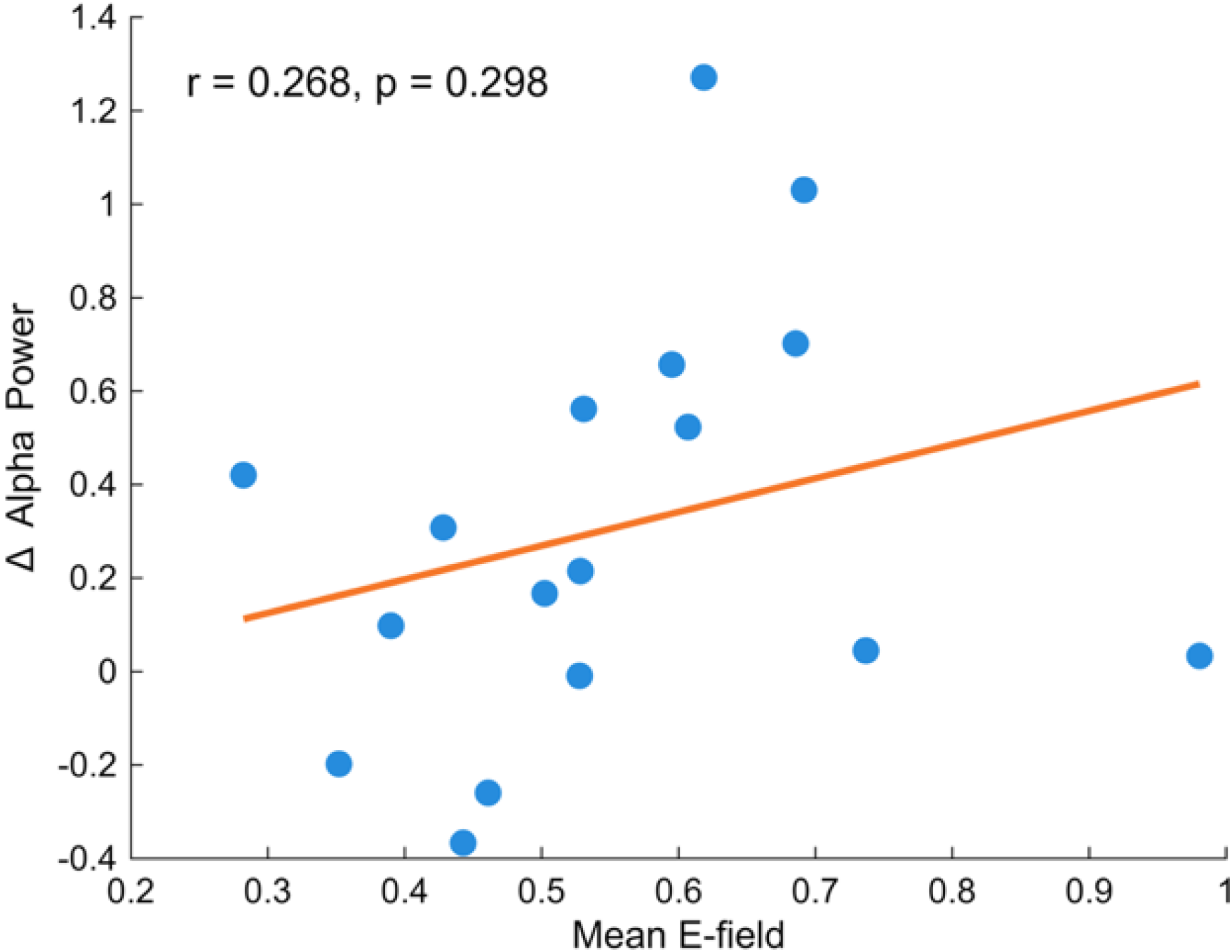
Correlation of individually simulated e-field strength and the relative change in alpha power between the pre- and post-measurements. Pearson correlation revealed no significant correlation (p=0.298), even though a visual trend is visible. Blue points depict individual data points; red line shows the regression line.

## Discussion

In this study we showed for the first time, that occipital tTIS is able to modulate occipital alpha power after 20 minutes of stimulation in comparison to sham stimulation and a high frequency carrier control while the latter two did not differ. However, this electrophysiological effect was not accompanied by behavioral changes, and the magnitude of alpha power change was not associated with the individually simulated e-field strength in the target region. Over all 51 stimulation sessions, the stimulation was very well tolerated, and no severe side effects were reported by the participants. These results suggest that the tTIS effect cannot be attributed to high-frequency stimulation or nonspecific session effects and there add valuable new knowledge which was unresolved in previous studies.^18,26^

The study conducted by von Conta et al. 2022 benchmarked if tTIS is able to reproduce tACS like effects in regard to occipital alpha power change after stimulation by comparing tTIS, tACS and an active high-frequency control.^18^ Interestingly, there were no significant difference between these stimulations which consequently lead to two possible explanations. One, that none of these conditions were able to produce a relevant alpha effect and second, that the control condition itself was physiologically active. These findings therefore produce an inconclusive three-condition pattern. As a consequence, the authors explicitly highlighted the need to further evaluate the effects of high carrier frequencies. This knowledge gap and the recommendation by von Conta et al. are specifically addressed in the present study by comparing tTIS, sham and a high-frequency carrier control condition to provide a more mechanistically interpretable framework. The almost absent difference between the carrier and the sham condition argues against physiologically effects of the carrier condition, i.e. being able to elicit alpha power changes. This favors the interpretation that the alpha increases reported in the earlier study may have reflected factors other than a carrier-specific stimulation effect. Our new findings extend the current knowledge by showing effects of tTIS behind those of sham or pure high frequency stimulation. Looking at the results of von Conta et al. they raised the question whether the carrier frequencies themselves have neuromodulatory effects and thus question a ground assumption of tTIS, i.e. that the overlaying cortex is not affected by the stimulation. In summary, the benchmark study of von Conta et al. established the feasibility and revealed a certain ambiguity of stimulation effects. To disentangle this ambiguity, the current study introduced sham and carrier control conditions and provided evidence for a tTIS specific stimulation effect which was not shown in the other two conditions.

Another fundamental question regarding the field is whether biological effects of tTIS are a result of the temporal interference stimulation, the high-frequency stimulation itself or an interaction of both.^26–28^ While originally introduced as membrane filtering based, later work showed that the underlying mechanism of tTIS seems to be more complex.^7^ Mirzakhalili et al., for example, argued for non-linear membrane processes and ion-channel-mediated rectifications are required to explain neurons sensitivity to TI-waveforms.^26^ Since carrier-only stimulation did not show a significant effect, our data strongly suggest that TI properties are necessary to elicit electrophysiological tTIS effects.

Occipital alpha power is one of the most extensively studied electrophysiological targets in the tACS field. Multiple studies demonstrated the ability of tACS to increase endogenous alpha power compared to sham stimulation.^29–31^ While the magnitude of effects varies across stimulation paradigms and investigated groups, they could be proven up to 70 minutes after actual stimulation.^32–34^ The current results are therefore placed in a well-established framework which has already proven that externally applied, sufficient electrical stimulation can modulate endogenous alpha oscillations. Our findings are well in line with this framework and suggest that this modulation, including aftereffects, is also possible by a temporal interference waveform at or near the individual alpha frequency. This is especially interesting since occipital alpha oscillations represent a proximal electrophysiological target rather than behavioral outcomes that rely on downstream processes. However, it is noteworthy that the results in our study reflect aftereffects of tTIS and thus are no demonstration of alpha entrainment during the stimulation. This is important since Vossen et al. showed that aftereffects lacked features expected from entrained oscillations and suggested plastic changes as the source of aftereffects.^35^ Similarly, Zaehle et al. discussed spike-time-dependent plasticity as a possible mechanism of tACS aftereffects. However, various potential mechanism could explain the results; one of them is an online entrainment followed by persisting aftereffects, another would be altered cortical excitability or synaptic plasticity. Given the current study design, the present study results cannot distinguish between these mechanisms and therefore should be interpreted as evidence for post-stimulation effect of tTIS rather than direct evidence for online neuronal entrainment.

Despite the significant alterations in the EEG data, the behavioral outcome did not show any effects of the stimulation which suggests a dissociation between electrophysiological markers of target engagement and behavioral performance. As behavior is a rather distal measure of neuromodulation, with multiple processing stages in between, this indicates that a visual vigilance task may not be sensitive to significant changes induced by electrical stimulation but does not invalidate the EEG findings. Furthermore, there is evidence that electrophysiological and behavioral outcomes are not necessarily congruent in electrical neuromodulation.^36^

Surprisingly, the correlation between the individually simulated e-field in the target region and the change in alpha power did not show a significant result. While Kasten et al. 2019 could show such a relation, the authors did incorporate more than only the e-field magnitude but also spatial correspondence.^37^ In addition, the sample size in this study may also account for these findings as it may have been too small to capture modest correlations. With regards to safety concerns and tolerability, we here provide further evidence that tTIS is a safe und feasible stimulation method which is in line with previous studies on these aspects. ^38^

In this article, we present a within-subject designed randomized, double-blinded and controlled study which minimizes interindividual variability and investigates direct physiological outcome for aftereffects of tTIS in young healthy subjects. On the other hand, some limitations also apply. With a small sample size of 17 participants, future studies should underpin our findings with larger study groups. In addition, we did not assess the online effects of tTIS and did not compare different stimulation patterns, e.g. continuous or intermittent theta burst. This limits the generalizability of our results in a certain way. To overcome these limitations, future studies should investigate larger sample sizes and test systematically the manipulation of different stimulation parameters like frequency, intensity or stimulation duration combined with online effects and suitable behavioral tasks.

In summary, we were able to significantly enhance occipital alpha power by using tTIS in comparison to both, a high-frequency carrier control and sham stimulation. This is the first study which proved the absence of significant carrier-/sham effects and is therefore able to attribute the effects to tTIS. Given the design of our study, this can be interpreted as electrophysiological aftereffects of temporal interference stimulation and therefore provides evidence that tTIS can selectively modulate neuronal oscillations and supports further investigation of different stimulation parameters.

## Data availability

The data that supports the findings in this manuscript are available upon resonable request.

